# CsFDL1-CsFTL3 complex represses *CsFTL3* via negative feedback to fine-tune flowering in *Chrysanthemum seticuspe*

**DOI:** 10.64898/2026.02.26.708152

**Authors:** Shuang Wang, Chunmeng Wang, Zishu Mei, Yiman Yang, Shijun Zhong, Jingyi Qiu, Zhenxing Wang, Likai Wang, Sumei Chen, Weimin Fang, Fadi Chen, Jiafu Jinag

**Affiliations:** State Key Laboratory of Crop Genetics and Germplasm Enhancement, Key Laboratory of Flower Biology and Germplasm Innovation, Key Laboratory of Landscaping, Ministry of Agriculture and Rural Affairs, College of Horticulture, Nanjing Agricultural University, Nanjing 211800, China; Zhongshan Biological Breeding Laboratory, Nanjing 210014, China; Sanya Institute of Nanjing Agricultural University, Sanya 572025, China

**Keywords:** Flowering time, Negative feedback loop, Transcriptional repression, CsFTL3-CsFDL1 complex, *Chrysanthemum*

## Abstract

In many flowering plants, the transition from vegetative growth to reproductive development is regulated by seasonal changes in photoperiod. Under inductive photoperiods, leaves produce the florigen FT (FLOWERING LOCUS T), which is transported to the shoot apex to promote flowering. The photoperiod is known to have a major effect on the flowering of chrysanthemum. In the perennial short-day (SD) plant *Chrysanthemum seticuspe*, the expression of *CsFTL3* (*FT-like* gene) does not increase immediately after shifting from long-day (LD) to SD conditions but gradually accumulates under continuous SD conditions, peaking during inflorescence development. However, the underlying mechanism remains elusive. We show that *CsFDL1* (an ortholog of FD) and *CsFTL3* exhibit a significant inverse expression pattern in leaves during the initial stage of short-day inductions. Furthermore, the expression of *CsFTL3* is upregulated in the leaves of *CsFDL1*-knockdown transgenic lines. CsFDL1 is expressed in leaves and forms a complex with CsFTL3 to recognize several TCGA- and ACGT-containing motifs in the *CsFTL3* promoter. The CsFTL3-CsFDL1 complex downregulates *CsFTL3* expression, thereby preventing its excessive induction by SD signals and inhibiting precocious floral transition. This study reveals that *CsFDL1* acts as a key early repressor in the photoperiodic flowering pathway of chrysanthemum leaf, mediating negative feedback regulation by forming a complex with CsFTL3 to achieve precise temporal control of short-day-dependent flowering responses.

## Introduction

Flowering represents a complex developmental process finely tuned by a multi-gene network, through the coordinated integration of environmental and endogenous cues (Freytes et al., 2021). In many plants, the transition from vegetative to reproductive growth is determined by seasonal changes in day length, a phenomenon known as photoperiod regulation (Song et al., 2015). The photoperiod pathway typically regulates the expression of a series of transcription factors through the endogenous circadian clock, ultimately inducing the production of the florigen gene *FT* to initiate floral transition (Andres and Coupland, 2012). The FT gene serves as a key integrator of multiple flowering pathways in leaves, converging signals from different pathways. Synthesized in leaves, FT protein is transported to the shoot apex, where it forms a complex with the bZIP transcription factor FD. This complex subsequently activates the expression of floral meristem identity genes such as *APETALA1* (*AP1*) and *FRUITFULL* (*FUL*), playing a central role in the floral transition (Wigge et al., 2005). This conserved regulatory mechanism of the FT-FD module has been confirmed in various plants, including *Arabidopsis thaliana* (Corbesier et al., 2007), Rice (Tamaki et al., 2007), and *Cucurbita moschata* (Lin et al., 2007). However, the *FT-like* gene family has evolved functionally diverse paralogous genes through gene duplication and functional divergence. These genes precisely regulate developmental processes such as flowering time and inflorescence architecture via antagonistic or synergistic interactions, adapting to different photoperiods and environmental conditions, thereby shaping diverse agronomic traits (Jin et al., 2021). For example, in rice, Hd3a and RFT1 coordinately regulate photoperiodic flowering and influence panicle branching (Izawa et al., 2002; Kojima et al., 2002); in onion, multiple *FT* paralogs antagonistically regulate flowering and bulb formation (Lee et al., 2013).

Chrysanthemum (*Chrysanthemum morifolium* Ramat.), as a typical short-day plant, exhibits high sensitivity to photoperiod signals in its flowering process. *C.* has emerged as a model plant species of cultivated chrysanthemums, especially for studies involving diploid and self-compatible pure lines (Gojo-0) (Sun et al., 2002). In chrysanthemum, several *FT* homologs have been identified. Among them, *CsFTL3* from *C. seticuspe* has been confirmed as a key florigen-encoding gene (Oda et al., 2012). Unlike *FT* in *Arabidopsis*, which is rapidly induced under inductive photoperiods (Ma et al., 2020; Liu et al., 2018), *CsFTL3* expression does not increase immediately after shifting from LD to SD conditions. Instead, it gradually accumulates under continuous SD treatment (Higuchi et al., 2013). This unique expression pattern suggests that *CsFTL3* is subject to a more complex transcriptional regulation tailored to the specific photoperiod requirements of chrysanthemum.

As a key interacting partner of FT, FD homologs play a crucial role in mediating FT-dependent flowering regulation (Taoka et al., 2011; Li et al., 2015). In *C. seticuspe*, *CsFDL1* is an FD-homologous bZIP transcription factor. Previous studies have shown that CsFDL1 interacts with the CsFTL3 protein (Higuchi et al., 2013; Nakano et al., 2019; Tian et al., 2025). However, the spatiotemporal expression pattern of *CsFDL1* during SD induction, its specific regulatory role in *CsFTL3* transcription, and whether the CsFDL1-CsFTL3 module possesses regulatory functions beyond activation during the early response to photoperiod signals remain unclear.

This study systematically analyzed the tissue-specific expression of *CsFTL3* and *CsFDL1* in *C. seticuspe* and their dynamic changes under SD induction. The interaction between CsFDL1 and CsFTL3 was validated both in vitro and in vivo through yeast two-hybrid and bimolecular fluorescence complementation assays. Chromatin immunoprecipitation, luciferase reporter assays, and yeast one-hybrid assays were employed to investigate the binding and regulatory activity of *CsFDL1* on the promoters of *CsFTL3* and its downstream flowering integrator gene *CsAFL1*. Furthermore, by constructing and analyzing *CsFDL1* knockdown transgenic lines, the biological functions of *CsFDL1* in regulating flowering time and plant architecture in chrysanthemum were elucidated. This study aims to clarify the regulatory mechanism underlying CsFDL1-mediated feedback on *CsFTL3* expression during the early stage of short-day induction in leaf, thereby providing new insights into the molecular basis of continuous short-day-dependent flowering in chrysanthemum.

## Materials and methods

### Plant material and growth conditions

*C. seticuspe* (Gojo-0) were provided by the Graduate School of Integrated Sciences for Life, Hiroshima University, Japan (Nakano et al., 2021). Plant materials were initially cultivated under LD conditions (16 h light/8 h dark, 23 °C, 75 % relative humidity) for approximately 45 days to prevent premature flowering, at which stage plants developed 14-16 fully expanded leaves. Subsequently, seedlings were transferred to a phytotron and exposed to SD conditions (8 h light/16 h dark, same temperature and humidity) to induce flowering (Cheng et al., 2023).

### RNA extraction and RT-qPCR analysis

Total RNA was extracted from each tissue using the Quick RNA Isolation kit (Huayueyang, Beijing, China). First-strand cDNA was synthesized from 1 μg RNA using the Evo M-MLV One Step RT-qPCR Kit (SYBR) (ACCURATE BIOTECHNOLOGY, Changsha, China). RT-qPCR was performed on a LightCycler 96 system (Roche, Basel, Switzerland) with the following program: 95 °C for 120 s, followed by 45 cycles of 95 °C for 15 s, 55 °C for 15 s, and 72 °C for 15 s. All reactions were performed in triplicate, both biological and technical. Gene expression levels were calculated using the 2^-ΔΔCT^ or 2^-ΔCT^ method (Hu et al., 2025), with CsACTIN as the reference gene (Higuchi et al., 2011). Primer sequences are listed in Supplementary Table S1.

### Yeast one-hybrid assay

For the yeast one-hybrid assay, the 1386 bp promoter region of *CsFTL3* was amplified and ligated into the Sac II-digested pHIS2 vector to serve as the bait. The full-length CDS of *CsFDL1* was cloned into the pGADT7 vector as the prey. Plasmids including pGADT7-*GUS*, pGADT7-*CsFDL1*, and pHis-*CsFTL3*pro were transformed into the yeast strain Y1H (Clontech, Mountain View, CA, USA), with pGADT7-*GUS* used as the negative control. Interactions were detected on synthetic defined medium lacking histidine, leucine, and tryptophan (SD/-His-Leu-Trp) supplemented with 3-amino-1,2,4-triazole (3-AT) at concentrations of 180 and 220 mM.

### Dual-luciferase reporter assay and protoplast transformation

The CDS of *CsFDL1* and *CsFTL3* was cloned into pORE-R4-35AA (Hu et al., 2025), and the 1386 bp *CsFTL3* promoter was ligated into *Spe* I-digested pGreenII 0800-LUC to generate the reporter plasmid. Two transformation assays were conducted: pORE-R4, pORE-R4-*CsFDL1*, pORE-R4-*CsFTL3*, and *CsFTL3*pro-0800-LUC were transformed into wild-type (WT) *C. seticuspe* protoplasts (Higuchi et al., 2013); all vectors were also transformed into *Agrobacterium tumefaciens* GV3101, with *CsFTL3*pro-0800-LUC co-transformed with pORE-R4-*CsFDL1*/ pORE-R4-*CsFTL3* into *Nicotiana benthamiana* leaves (pORE-R4-35SAA as negative control). LUC/REN ratios were measured using a Dual-Luciferase Reporter Gene Assay kit (Yeasen, Shanghai, China), and LUC activity was detected using a CCD imaging system (Tanon 5200, Shanghai, China).

### Chromatin immunoprecipitation-qPCR assay

The tobacco rattle virus (TRV)-derived engineered vectors pTRV1 and pTRV2 were used to construct the transient overexpression vector OE-TRV2 (Huang et al., 2022). OE-TRV2-*CsFDL1*-HA transgenic plants were generated (Supplementary Fig. S2), and both these transgenic plants and WT plants were subjected to chromatin immunoprecipitation-quantitative PCR (ChIP–qPCR) analysis. Pierce™ ChIP-grade Protein A/G Magnetic Beads (Thermo Fisher Scientific, Waltham, MA, USA) and HA recombinant rabbit monoclonal antibodies (Thermo Fisher Scientific) were employed to enrich target DNA fragments. Subsequently, the enriched DNA fragments were detected by reverse transcription-quantitative PCR (RT-qPCR) using the primer pairs listed in Supplementary Table S1.

### Yeast two-hybrid assay

The coding sequences of *CsFTL3* and *CsFDL1* were cloned into the pGADT7 and pGBKT7 vectors. The paired recombinant plasmids were transformed into the yeast strain *Saccharomyces cerevisiae* Y2H and selected on synthetic defined medium lacking leucine and tryptophan (SD/-Leu-Trp). The pGBK-53/pGAD-T combination served as a positive control, and pGBK-Lam/pGAD-T as a negative control. Transformants were incubated at 28 °C for 3 days on SD/-Leu-Trp medium, then replica-plated onto quadruple dropout medium (SD/-Leu-Trp-His-Ade). Positive clones were identified using 5-bromo-4-chloro-3-indolyl-α-D-galactopyranoside (X-α-Gal) screening. The primer pairs are listed in Supplementary Table S1.

### BiFC assay

The coding sequences of *CsFTL3* and *CsFDL1* were cloned into pSPYNE and pSPYCE. The recombinant plasmid combinations pSPYNE-*CsFTL3* + pSPYCE-*CsFDL1*, pSPYNE-*CsFTL3* + pSPYCE, and pSPYNE+pSPYCE-*CsFDL1* were separately introduced into *A. tumefaciens* strain GV3101. The resulting bacterial suspensions were infiltrated into tobacco leaves. Following 24 h of dark incubation and 24 h of light incubation, yellow fluorescent protein (YFP) and red fluorescent protein (RFP) signals were observed using a laser-scanning confocal microscope (Zeiss LSM800, Germany).

### Luciferase complementation (LCI) assay

The open reading frames (ORFs) of *CsFTL3* and *CsFDL1* were cloned into the pCAMBIA1300-nLUC (nLUC) and pCAMBIA1300-cLUC (cLUC) vectors, respectively, and then introduced into *A. tumefaciens* strain GV3101. The transformed *A. tumefaciens* was resuspended in infiltration buffer and injected into leaves of 5-week-old *N. benthamiana* plants. After 24 h of dark incubation followed by 48 h under LD conditions, 100 mM sodium fluorescein salt was sprayed onto the leaves, which were then kept in darkness for 5 min. Luciferase (LUC) activity was detected, and images were captured using a CCD imaging system (Tanon 5200, Shanghai, China), as previously described.

### Plant transformation and phenotype analysis

To construct the knockdown vector pORE-R4-amiR-*CsFDL1*, four oligonucleotides (oligos) were designed and synthesized using Web MicroRNA Designer (https://wmd3.weigelworld.org/cgi-bin/webapp.cgi). The pORE-R4 vector was then constructed with Sal I and Spe I as restriction enzymes. For plant transformation, the knockdown plasmid (pORE-R4-amiR-*CsFDL1*) was introduced into *A. tumefaciens* strain EHA105, and transgenic chrysanthemum plants were obtained via the *A. tumefaciens-mediated* leaf-disc infection method, as described by Li et al. (2015). Primer pairs 35S-F/II (Table S1) were designed to validate the transgenic lines at the DNA level, and oligos qRT-*CsFDL1*-F/R (Table S1) were used for further confirmation of positive transgenic lines. Wild-type and transgenic plants were grown at 23°C with LD (16 h light/8 h dark) and SD (8 h light/16 h dark) conditions. The time was recorded when the plant first showed visible flower buds. For each strain, 12 plants were analyzed. Significant differences between groups were determined using DPS_7.05 software (Tang et al., 2012).

## Results

### Expression patterns of *CsFTL3* and *CsFDL1* in C. seticuspe

To investigate the expression patterns of *CsFTL3* and *CsFDL1*, we harvested tissues from *C. seticuspe* at both vegetative and reproductive growth stages for RT-qPCR analysis. The results showed that *CsFTL3* was highly expressed in leaves during the reproductive stage (Fig. 1B). In contrast, *CsFDL1* was broadly expressed in leaves and shoot tips, with the highest transcript levels detected in roots (Fig. 1A). To further clarify the roles of *CsFTL3* and *CsFDL1* in photoperiod-dependent flowering regulation, we analyzed their dynamic expression patterns in leaves after transferring plants from LD to inductive SD conditions. The absolute expression level of *CsFTL3* was very low under LD conditions. There was a significantly downregulated within the first week after the shift to SD conditions, then gradually recovered and continued to increase (Fig. 1D). In contrast, *CsFDL1* was specifically induced in the first week of SD treatment, and then gradually decrease (Fig. 1C). Moreover, *CsFTL3* and *CsFDL1* transcript levels were detected in shoot tips, there were no remarkable differences in *CsFTL3* and *CsFDL1* expression during the first week after shifting from LD to SD condition (Supplementary Fig. S1). Subsequently, *CsFDL1* expression gradually decreased, whereas *CsFTL3* was progressively upregulated.

**Fig.1.**
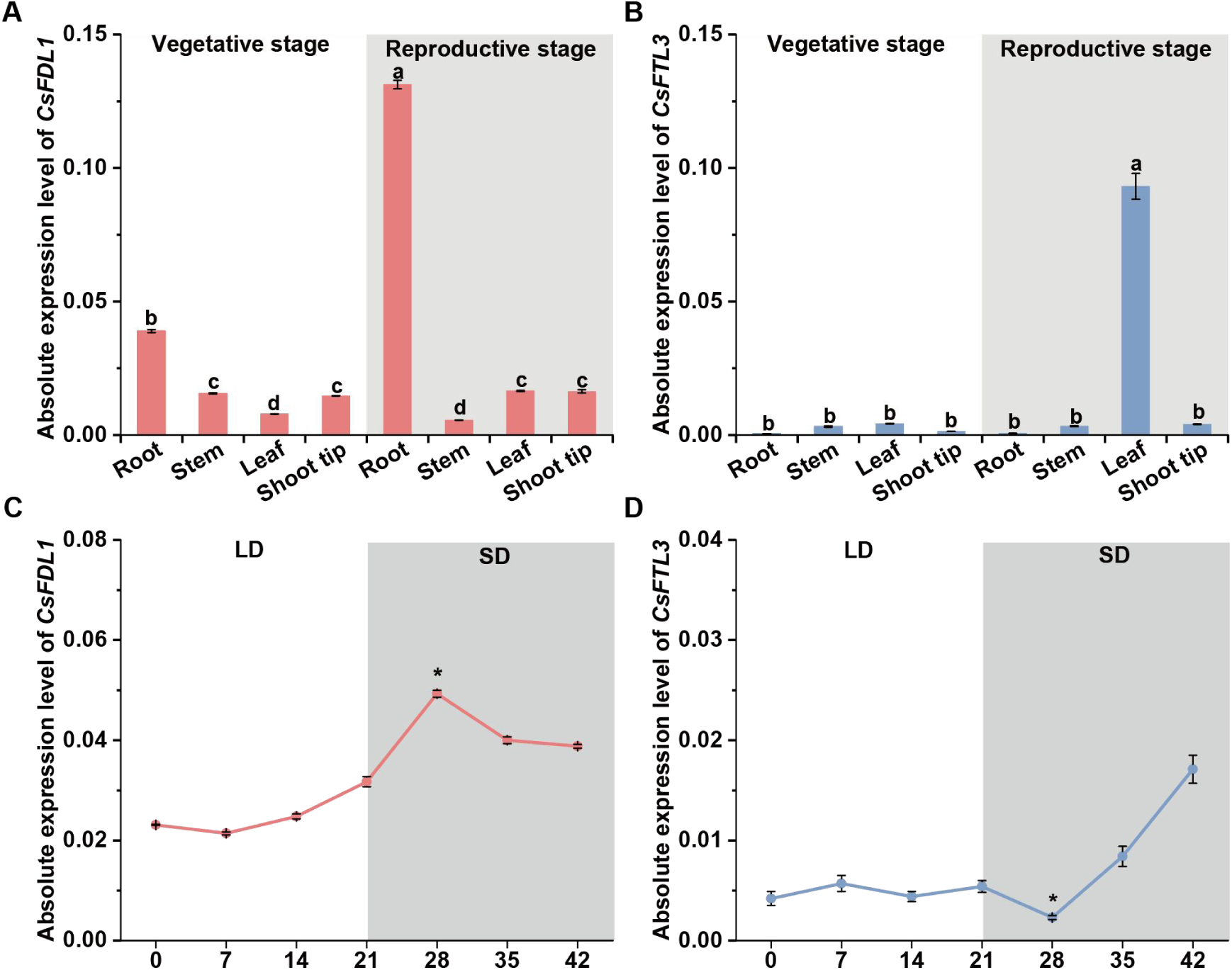
Expression pattern analysis of *CsFTL3* and *CsFDL1* in *C. seticuspe*. Transcript levels of *CsFDL1* (A) and *CsFTL3* (B) in different tissues at the vegetative and reproductive stages. Letters above the bars indicate significant differences as determined by Tukey’s test (P<0.05). Dynamic expression of *CsFDL1* (C) and *CsFTL3* (D) in leaves at different time points after transfer from LD (white background) to SD (gray background) conditions. Error bars indicate ±SD; n≥9. **P<0.01, *P<0.05 (Student’s t-test).

### *CsFDL1* regulates flowering

To investigate the role of *CsFDL1* in the floral transition of *C. seticuspe*, *CsFDL1* knockdown transgenic lines were generated using an artificial microRNA (amiRNA). Following PCR confirmation at the DNA level, four independent knockdown lines were obtained (Supplementary Fig. S2A). Two lines were randomly selected for phenotypic analysis. Knockdown of *CsFDL1* expression in *C. seticuspe* resulted in extremely late flowering under SD conditions (Fig. 2A, B, E). There were no remarkable differences in leaf number in WT and amiR-*CsFDL1* lines when WT plants showed visible buds (Fig. 2D). However, the amiR-*CsFDL1* plants exhibited shorter internodes and reduced plant height (Fig. 2C), suggesting that *CsFDL1* may also function in regulating plant architecture.

**Fig.2.**
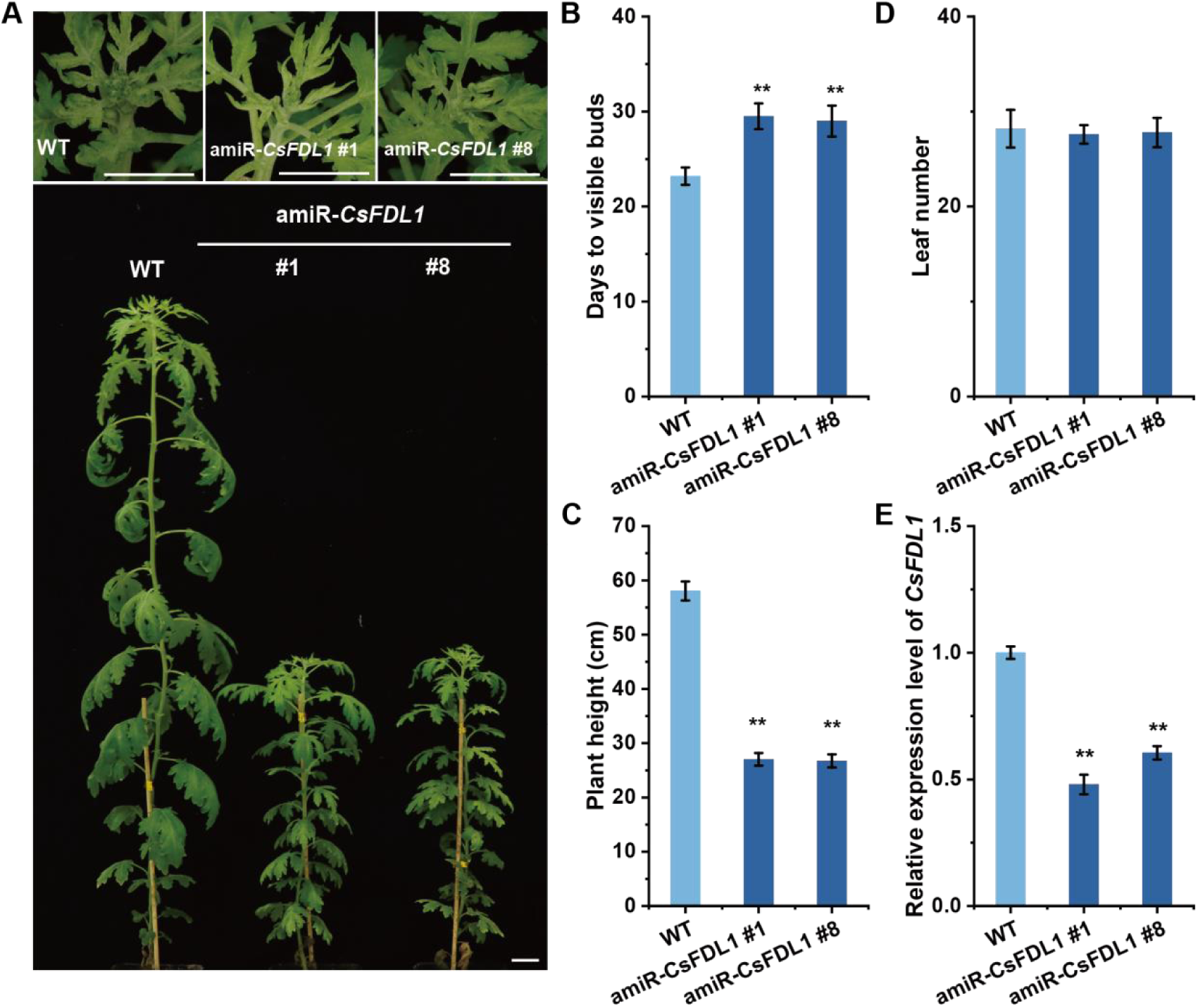
The phenotype of *CsFDL1* knockdown plants under SD conditions. (A)The phenotype of amiR-*CsFDL1* transgenic plants and WT at the bud stage, bar = 3 cm; (B, C, D) Statistics of flower bud emergence time, plant height, and leaf number in WT and amiR-*CsFDL1* transgenic plants. Error bars indicate standard deviation (SD); n≥12. (E) Validation of the expression level of *CsFDL1* in transgenic lines and WT with qRT-PCR. amiR-*CsFDL1* #1 and amiR-*CsFDL1* #8 represent the two independent *CsFDL1* knock-down lines; WT: wild-type plant. Values are mean±SE (n = 3); **P<0.01, *P<0.05 (Student’s t-test).

### *CsAFL1* and *CsFTL3* were repressed in amiR-*CsFDL1* plants

The FT–FD protein complex triggers a cascade of positive transcriptional events during floral induction, including the activation of CsAFL1 (an AP1/FUL-like gene) and CsM111 (an AP1 homolog) (Taoka et al., 2011; Higuchi et al., 2013). To further explore how *CsFDL1* regulates downstream flowering genes at SD, we examined the transcript levels of *CsFTL3*, *CsAFL1*, and *CsSOC1* in leaves of amiR-*CsFDL1* plants at 0, 4, and 8 days of SD induction. *CsFTL3* is highly expressed in the *CsFDL1* knockdown plant’s leaves in the leaves under SD conditions (Fig. 3A). Similarly, *CsAFL1* expression was significantly elevated at 0 and 4 days, but no significant difference was observed at 8 days (Fig. 3C). To elucidate the regulatory mechanism of *CsFDL1* on *CsAFL1*, yeast one-hybrid assays confirmed that *CsFDL1* directly binds to the promoter region of *CsAFL1*, indicating that *CsFDL1* exerts its inhibitory effect on *CsAFL1* through direct transcriptional regulation (Supplementary Fig. S6). Further examination of gene expression in shoot apices and leaves after 12 days of SD induction showed the expression level of *CsFTL3* in the WT was higher than that in the *CsFDL1* knockdown lines, whereas *CsAFL1* expression was significantly reduced in the knockdown lines (Fig. 3B, D) (*CsSOC1* expression is detailed in Supplementary Fig. S3). These findings suggest that during the later stages of SD induction, long-range feedback signals originating from the shoot apex may dominate *CsAFL1* expression in leaves, leading to a gradual convergence of expression differences between *CsFDL1* knockdown lines and the WT.

**Fig.3.**
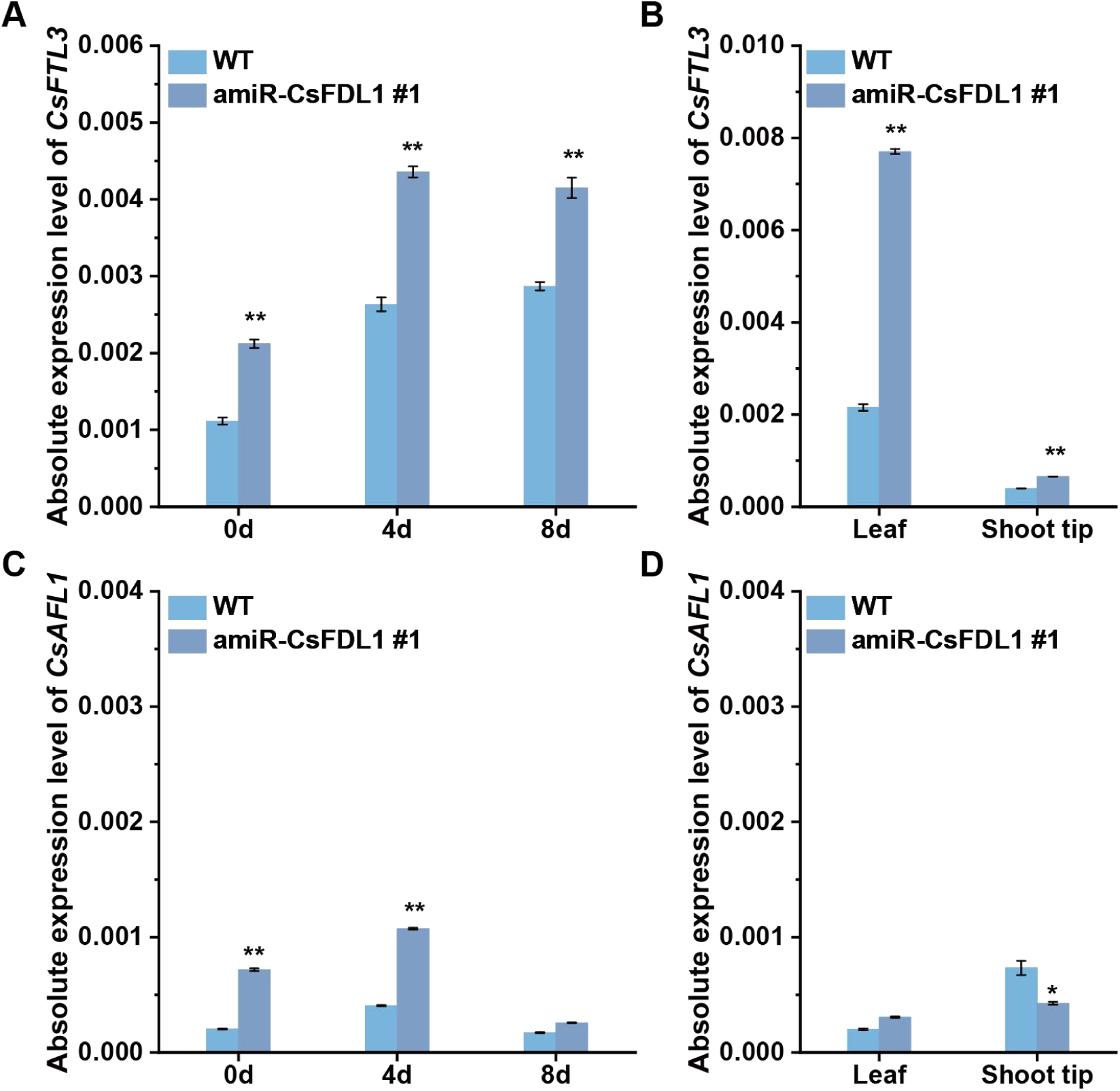
Relative expression levels of *CsFTL3* and *CsAFL1* in WT and amiR-*CsFDL1* plants under SD induction. (A, C) Expression of *CsFTL3* and *CsAFL1* was analyzed by qRT-PCR at 0, 4, and 8 days after SD induction (8 h light/16 h dark). (B, D) Expression of *CsFTL3* and *CsAFL1* in leaves and shoot tips was examined by qRT-PCR at 12 days after SD induction. Gene expression levels were calculated using the 2^-ΔCT^ method. Error bars represent standard deviation (SD); n = 3 independent experiments. **P < 0.01, *P < 0.05 (Student’s t-test).

### Coexpression with *CsFDL1* suppressed the autoregulation of *CsFTL3*

*CsFDL1* and *CsFTL3* exhibited opposite expression patterns under SD conditions. To evaluate the regulation, we conducted a transient gene expression experiment in protoplasts derived from mesophyll cells of *C. seticusp*e leaves. Expression of endogenous *CsFTL3* was up-regulated when *CsFTL3* was expressed alone or coexpressed with *CsFDL1* in WT protoplasts (Fig. 4). It has been shown that the CsFTL3-CsFDL1 complex establishes a photoperiod-dependent positive feedback loop in leaves, progressively amplifying the florigen signal (Higuchi et al., 2013). However, coexpression of *CsFDL1* and *CsFTL3* resulted in significant downregulation of endogenous *CsFTL3* transcript levels in WT protoplasts. In contrast, only a modest and statistically insignificant reduction was detected in *CsFDL1* knockdown transgenic *C. seticusp*e protoplasts (Fig. 4). These findings suggest that, while the CsFTL3-CsFDL1 complex drives a photoperiod-dependent positive feedback loop to amplify florigen signaling at the systemic level, its molecular regulatory mechanism is intricate and potentially conditional. CsFDL1 not only functions as an interacting partner of CsFTL3 to activate downstream genes but may also exert negative regulatory effects on *CsFTL3* transcription under specific conditions, such as particular cellular environments or expression levels. This could represent a mechanism for achieving homeostatic control or timely termination of the feedback loop.

**Fig.4.**
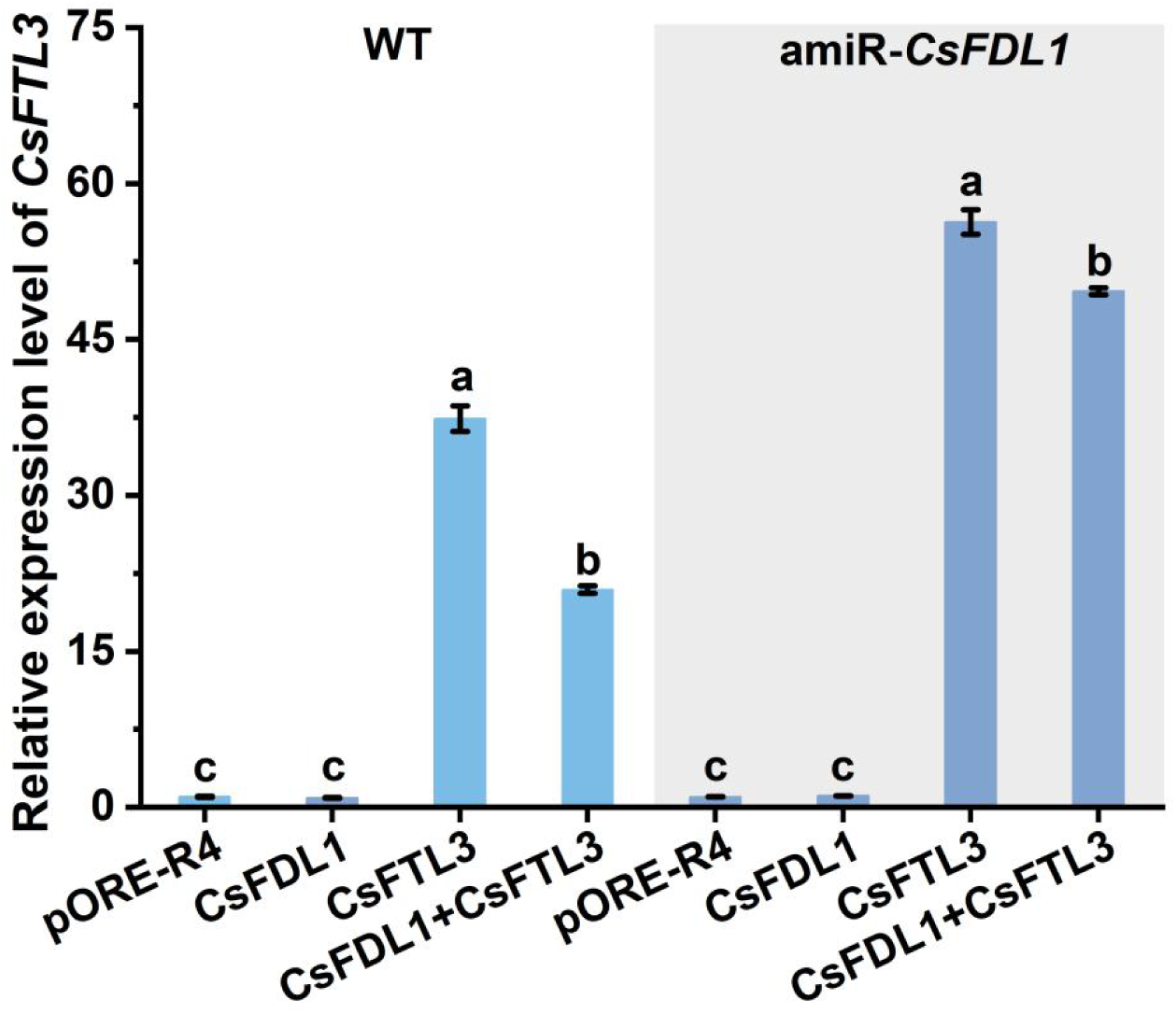
Regulation of *CsFTL3* by *CsFDL1* and *CsFTL3* in *C. seticuspe* protoplasts. (A) WT protoplasts; (B) amiR-CsFDL1#1 protoplasts. Transient expression assays were conducted in protoplasts isolated from WT and amiR-CsFDL1 #1 plants using different effector constructs (pORE-R4 as an empty vector control, CsFDL1, CsFTL3, and CsFDL1+CsFTL3). Error bars represent the mean±SD of three biological replicates. Different letters above the bars indicated significant differences based on Tukey’s HSD test (p < 0.05).

### CsFDL1-CsFTL3 complex formation in vivo

It is reported that FT acts as a transcriptional regulator and activates the expression of downstream flowering genes by forming the FT-FD-14-3-3-DNA complex (Gao et al., 2025). To explore the interaction between CsFDL1 and CsFTL3, BD-CsFDL1 and AD-CsFTL3 vectors were constructed for yeast two-hybrid assays. The results indicated no direct physical interaction between the two proteins (Supplementary Fig. S4), which is consistent with a previous report (Higuchi et al., 2013). We further used the tobacco (*Nicotiana benthamiana*) co-expression system to perform bimolecular fluorescence complementation (BiFC) assays (Fig. 5A) and firefly luciferase complementation imaging (LCI) assays (Fig. 5B). Both assays validated the in vivo interaction between CsFDL1 and CsFTL3, demonstrating their ability to form heterocomplexes. This result indicates that CsFDL1 and CsFTL3 can function in the same transcriptional complex.

**Fig.5.**
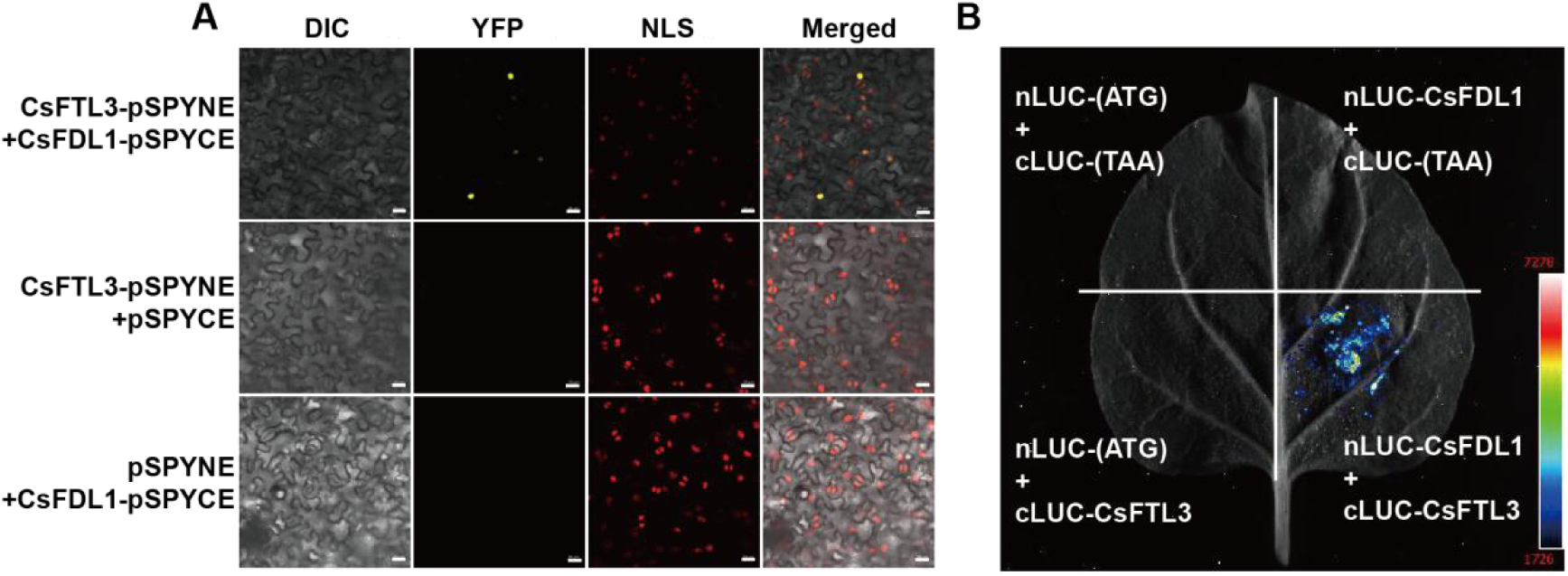
CsFDL1 interacts with CsFTL3 in vivo. (A) Bimolecular fluorescence complementation (BiFC) assay showing the interaction between CsFDL1 and CsFTL3. D53-mCherry, a nuclear-localized marker (NLS-mCherry), was used to indicate nuclear position. The pSPYNE and pSPYCE empty vectors served as negative controls. Scale Bars:20μm; (B) Firefly luciferase complementation imaging (LCI) assay confirming the interaction between CsFDL1 and CsFTL3.

### The CsFDL1-CsFTL3 complex represses *CsFTL3*

Recent studies suggest that the FT-FD complex can recognize and bind to ACGT or TCGA-containing motifs on the *FT* promoter, thereby inhibiting promoter cyclization and antagonizing the transcriptional activation of FT mediated by the CO-NF-Y complex (Abe et al., 2005; Collani et al., 2019). We identified the promoters of *CsFTL3* in the Chrysanthemum Genome Database (http://210.22.121.250:8880/asteraceae/homePage) and searched for ACGT and TCGA-containing motifs. TCGA and ACGT-containing motifs were identified near the distal CCAAT enhancer in the promoter region of *CsFTL3* (Fig. 6B). Subsequently, we performed a yeast one-hybrid (Y1H) assay, which showed that CsFDL1 proteins interacted with the promoters of *CsFTL3* (Fig. 6A). To test the CsFDL1 binding region of the *CsFTL3* genome sequences, we conducted a ChIP-qPCR experiment using OE-TRV2-*CsFDL1*:HA plants (Supplementary Fig. S2B, C). The P4 fragment served as controls in the 3’UTR regions of CsFTL3 (Fig. 6B). We found that CsFDL1 exhibited specific enrichment in the P2 region of the *CsFTL3* promoter (Fig. 6C).

**Fig.6.**
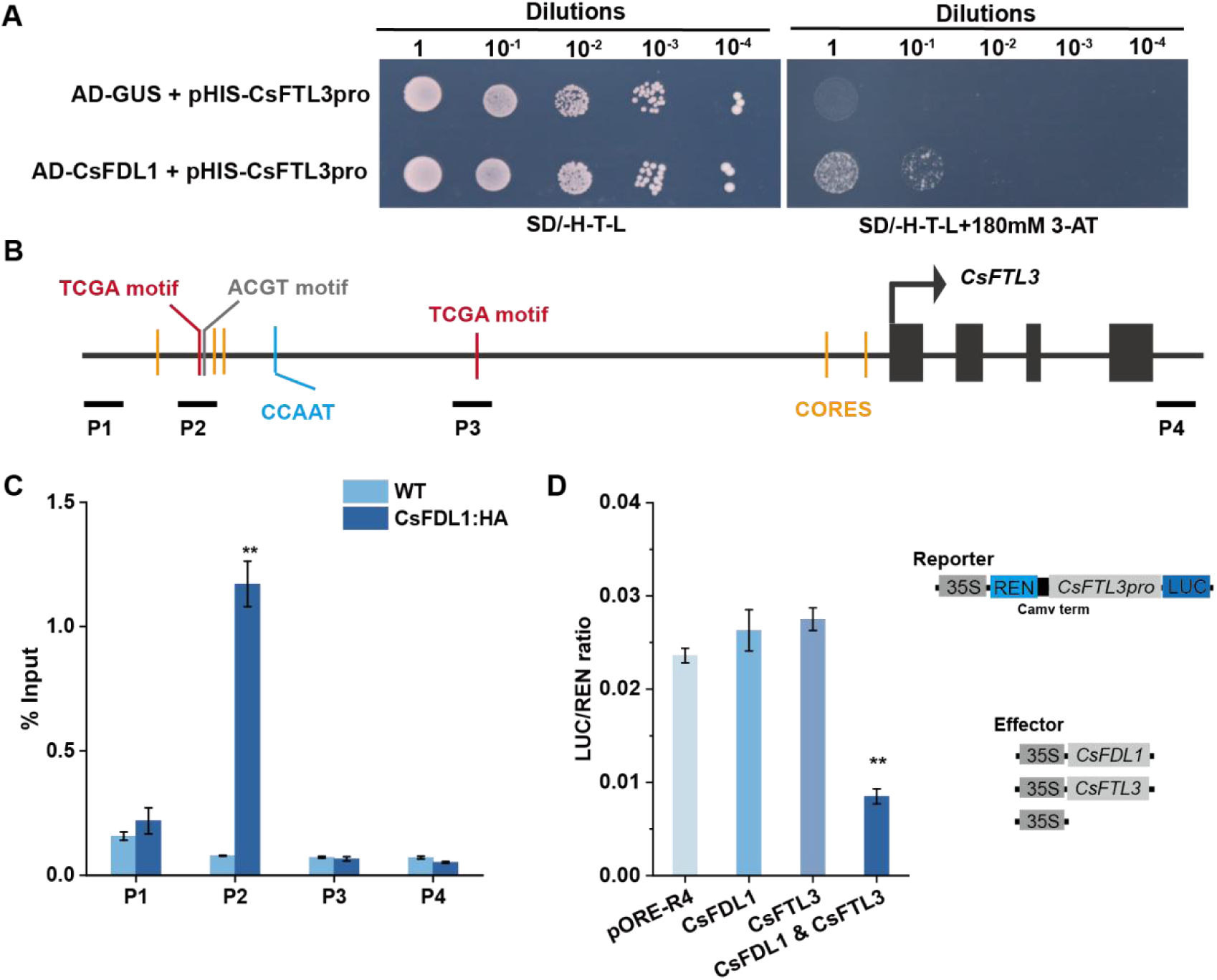
CsFDL1-CsFTL3 complex directly binds to the *CsFTL3* promoter and inhibits its transcription. (A) Yeast one-hybrid assay of the binding of *CsFDL1* to the promoter of *CsFTL3*; (B) Structure and fragments of the *CsFTL3* promoter used for ChIP-qPCR analysis. P1-P4: A variety of promoter segments were examined by RT-qPCR; (C) ChIP-qPCR assays of the regulatory regions of downstream genes from OE-TRV2-*CsFDL1*-HA transgenic plants. The data shown are presented as mean values with standard errors; n=3; **P<0.01 (Student’s t-test). (D) LUC/REN ratio represents the relative activity of the different effectors targeted with *CsFTL3*pro in chrysanthemum protoplast. The data are shown as the mean ± SE (n=3); **P<0.01(Student’s t-test).

To evaluate the effect of CsFDL1 on the regulation of *CsFTL3* promoter, a dual-luciferase reporter assay was selected for analysis. We observed that luciferase (LUC) activities derived from *CsFTL3* was significantly reduced when *CsFDL1* and *CsFTL3* coexpressed (Supplementary Fig. S5). Moreover, a luciferase (LUC) reporter system driven by the *CsFTL3* promoter was used to analyse *CsFTL3* expression. Compared with the empty vector control, neither *CsFDL1* nor *CsFTL3* alone could affect LUC activity driven by the *CsFTL*3 promoter. However, co-expression of *CsFDL1* and *CsFTL3* resulted in a significant reduction in LUC activity (Fig. 6D). Thus, we demonstrated a mechanism by which, the CsFDL1-CsFTL3 complex inhibits the transcription of *CsFTL3* in chrysanthemum by binding to ACGT- or TCGA-containing motifs.

## Discussion

The photoperiodic response in plants is a highly complex and precisely regulated process that involves the functional diversification and network reorganization of conserved regulatory modules across species. In hybrid aspen, the *FD* homologous gene has evolved dual functions: *FDL1* forms a complex with the FT2 protein to regulate SD-induced growth cessation, while also independently interacting with the transcription factor ABI3 in the abscisic acid signaling pathway to directly activate the expression of adaptive genes associated with stress resistance and bud maturation (Tylewicz et al., 2015). This indicates that *FDL1* serves as a core integrative node, synchronizing growth cycles with seasonal adaptive responses by switching interaction partners under different photoperiodic conditions, thereby highlighting the central role and evolutionary plasticity of the core components of the “FD-FT” module in plant environmental adaptation. In *Arabidopsis*, structural and biochemical analyses by Lv et al. (2021) elucidated the molecular mechanism by which the key transcription factor CO (CONSTANS) forms a heterotrimeric complex with NF-YB/YC and precisely regulates the expression of the florigen gene *FT* through multivalent binding. Notably, the FD-FT protein complex can suppress *FT* expression by interfering with the interaction between CO and NF-YB/YC, further underscoring the complexity and hierarchical nature of this regulatory network (Tian et al., 2025).

Previous studies have demonstrated that CsFTL3 and CsFDL1 can form a transcriptional activation complex under continuous SD conditions, promoting *CsFTL3* expression via a positive feedback loop, as elucidated by Higuchi et al. (2013), Nakano et al. (2019), and Tian et al. (2025). However, our spatiotemporal expression profiling revealed that during the early phase of SD induction, *CsFDL1* and *CsFTL3* exhibit opposing expression patterns in leaves: *CsFDL1* is rapidly induced as an early SD-responsive factor, whereas *CsFTL3* expression is significantly suppressed. The gradual accumulation of *CsFTL3* may reflect the chrysanthemum’s “memory” of continuous SD exposure, with the rapid induction of *CsFDL1* acting as an SD signal “sensor”, and the delayed expression of *CsFTL3* serving as a “verification mechanism” to ensure that irreversible inflorescence development is initiated only under persistent and stable SD conditions.

Thus, our current work provides a novel molecular framework in which *CsFDL1* acts as an early responsive factor to SD signals. CsFDL1 does not directly suppress the basal expression of *CsFTL3* but antagonizes its autoregulatory effect in a dose-dependent manner, confirming the existence of a negative feedback regulatory loop between the two genes (Fig. 7). This study advances the molecular mechanism of chrysanthemum floral induction from a “simple activation model” to a dynamic equilibrium model, where the precise temporal balance between activating and inhibitory signals determines the accuracy of the floral transition. This finding provides a new theoretical perspective for understanding how plants integrate environmental signals to regulate developmental timing. Future research should focus on elucidating the interactomes and regulatory networks of *FD* homologs across different species, thereby providing a more comprehensive understanding of the molecular basis of plant developmental plasticity and environmental adaptability.

**Fig.7.**
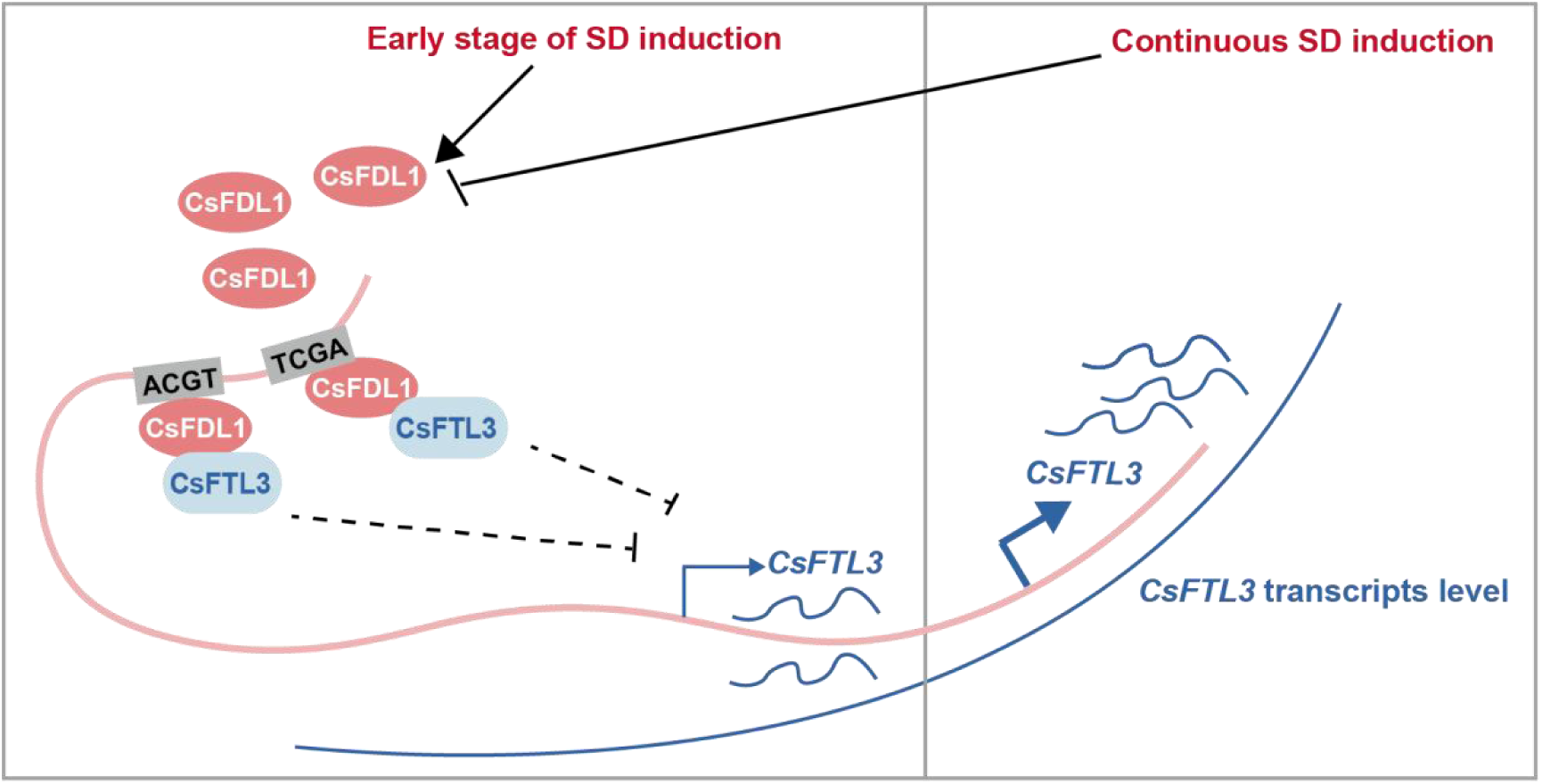
A working model for the dynamic equilibrium of the CsFTL3-CsFDL1 module in photoperiodic flowering.

## Acknowledgements

This work was financially supported by grants from the National Natural Science Foundation of China (32430096, 32272756), Zhongshan Laboratory for Biological Breeding Project (ZSBBL-KY2023-08), and a project funded by the Priority Academic Program Development of Jiangsu Higher Education Institutions. We thank Dr. Yuehua Ma (Central Laboratory of College of Horticulture, Nanjing Agricultural University) for assistance in using the laser scanning confocal microscope.

## Author contributions

JJ conceived and designed the experiments; SW performed most of the experiments; CW, ZM, YY, SZ and JQ provided technical support; ZW, LW and WF provided conceptual advice; SW and JJ analysed the data and wrote the manuscript; and SC and FC edited the manuscript.

## Supplementary Data

**Fig.S1.**
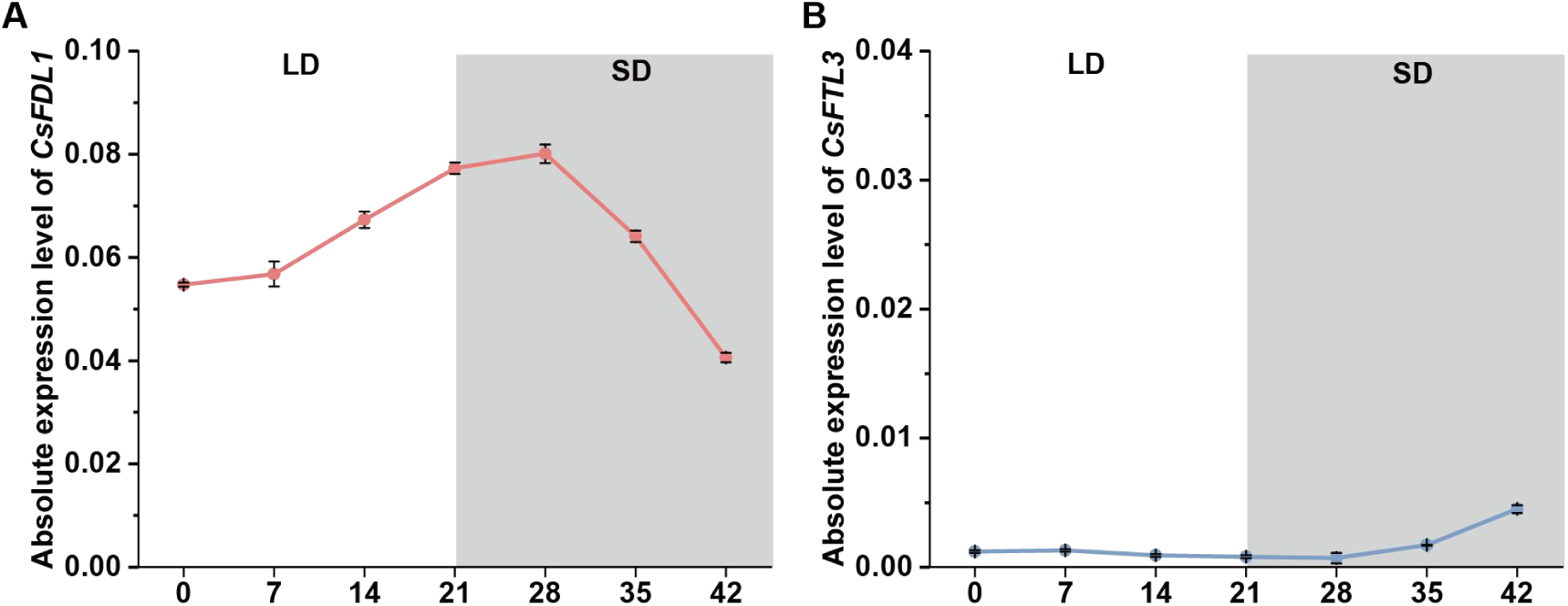
Dynamic expression of *CsFDL1* and *CsFTL3* in shoot tips at different time points after transfer from LD (white background) to SD (gray background) conditions. Error bars indicate ±SD; n≥9.

**Fig.S2.**
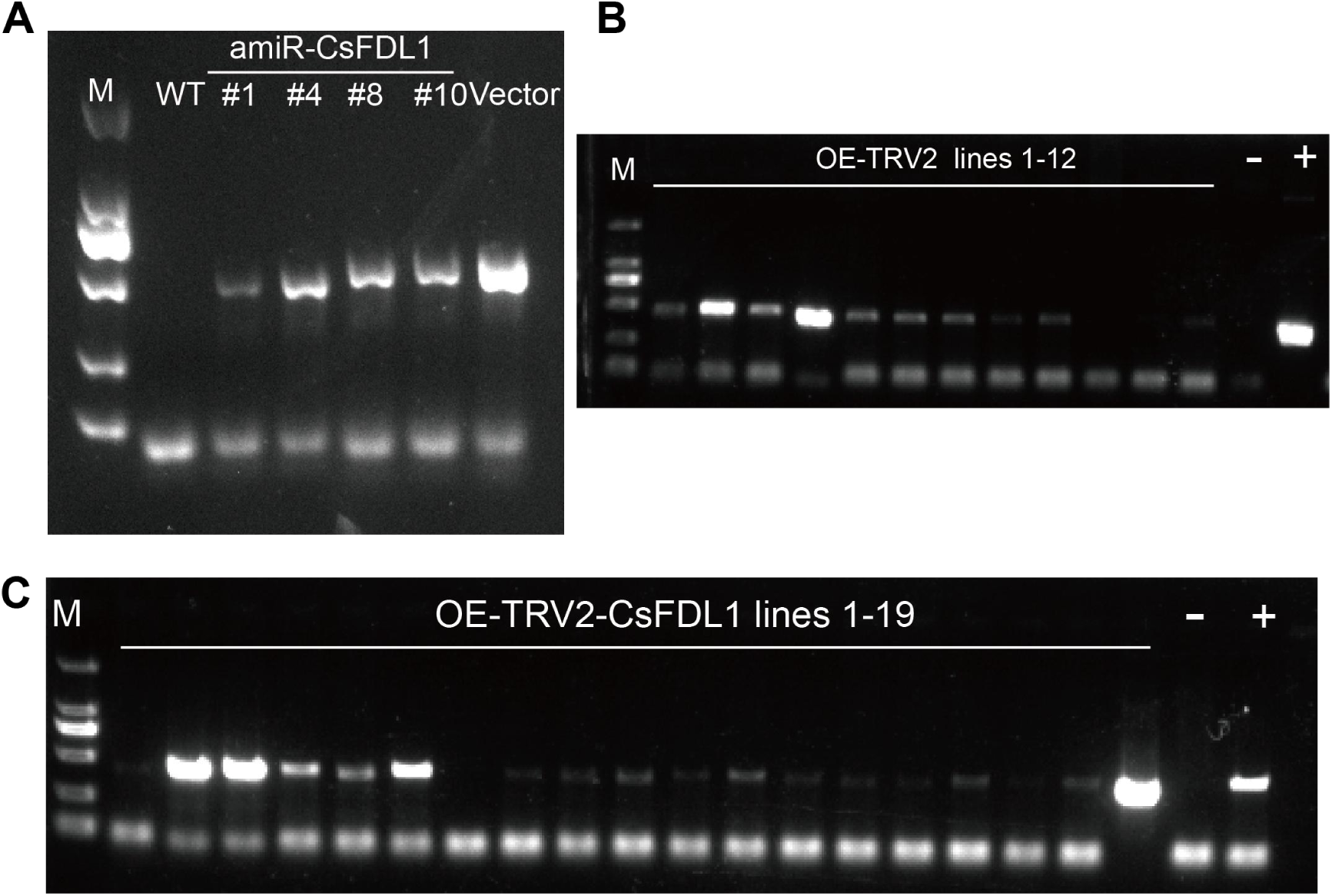
Identification of *CsFDL1* transgenic lines. Identification of *CsFDL1* transgenic lines via PCR at the DNA level. M: DL2000; Vector: Positive control; WT/-: Wild-type *C. seticuspe*.

**Fig.S3.**
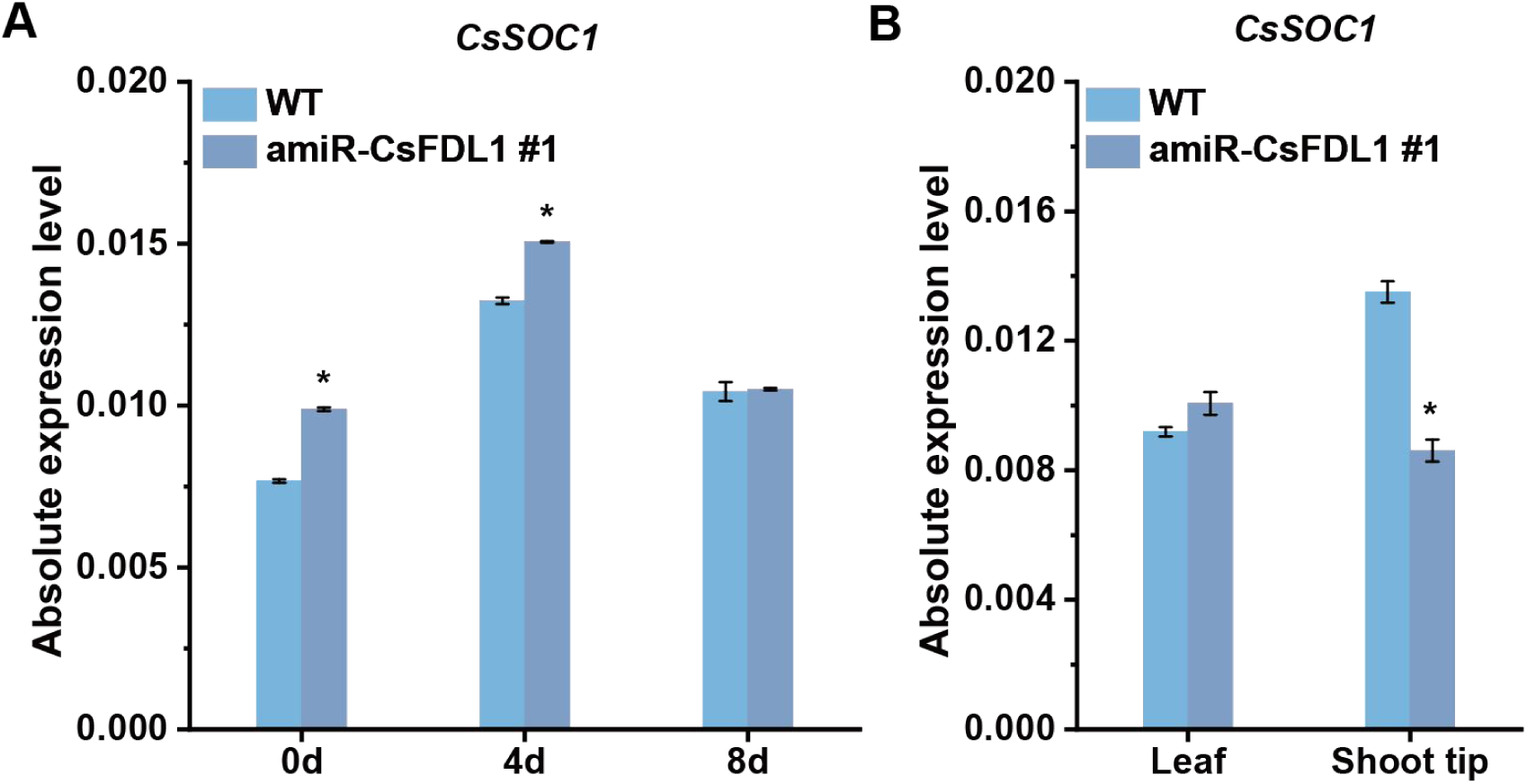
Relative expression levels of CsSOC1 in WT and amiR-CsFDL1 plants under SD induction. (A)Expression of CsSOC1 was analyzed by qRT-PCR at 0, 4, and 8 days after SD induction (8 h light/16 h dark). (B) Expression of CsSOC1 in leaves and shoot tips was examined by qRT-PCR at 12 days after SD induction. Gene expression levels were calculated using the 2^-ΔCT^ method. Error bars represent standard deviation (SD); n = 3 independent experiments. *P < 0.05 (Student’s t-test).

**Fig.S4.**
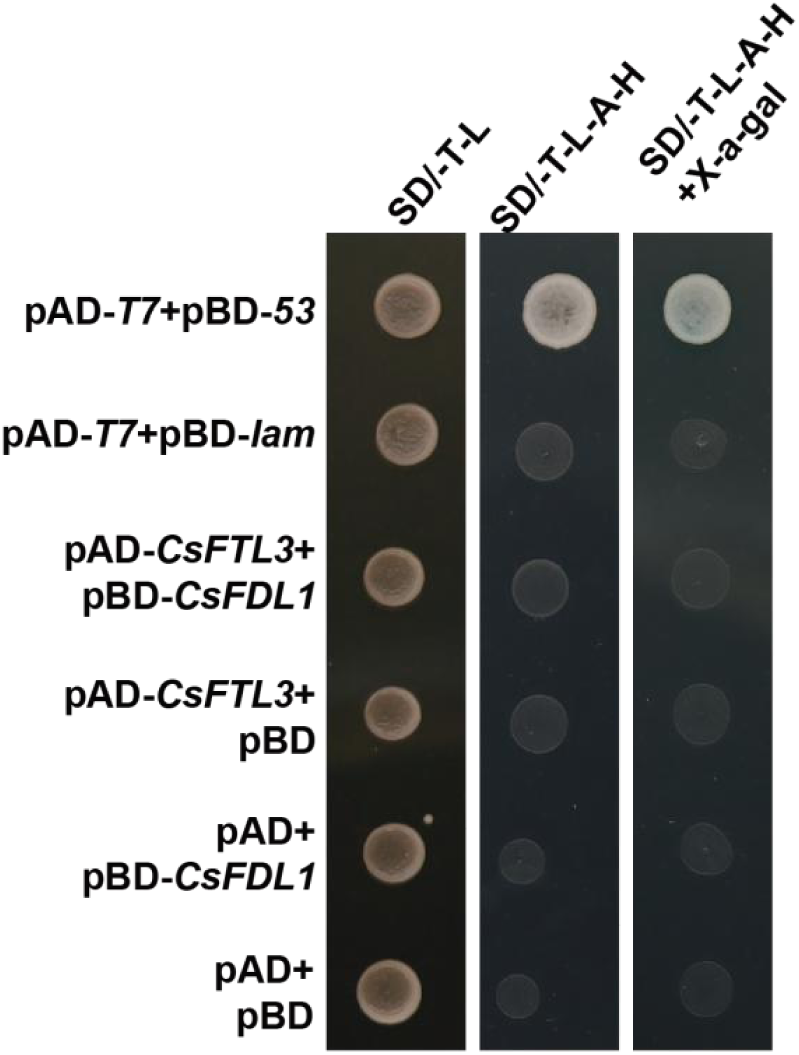
The interaction between CsFTL3 and CsFDL1 in yeast. Positive: pGBKT7-53& pGAD-T; Negative: pGBKT7-Lam& pGAD-T.

**Fig.S5.**
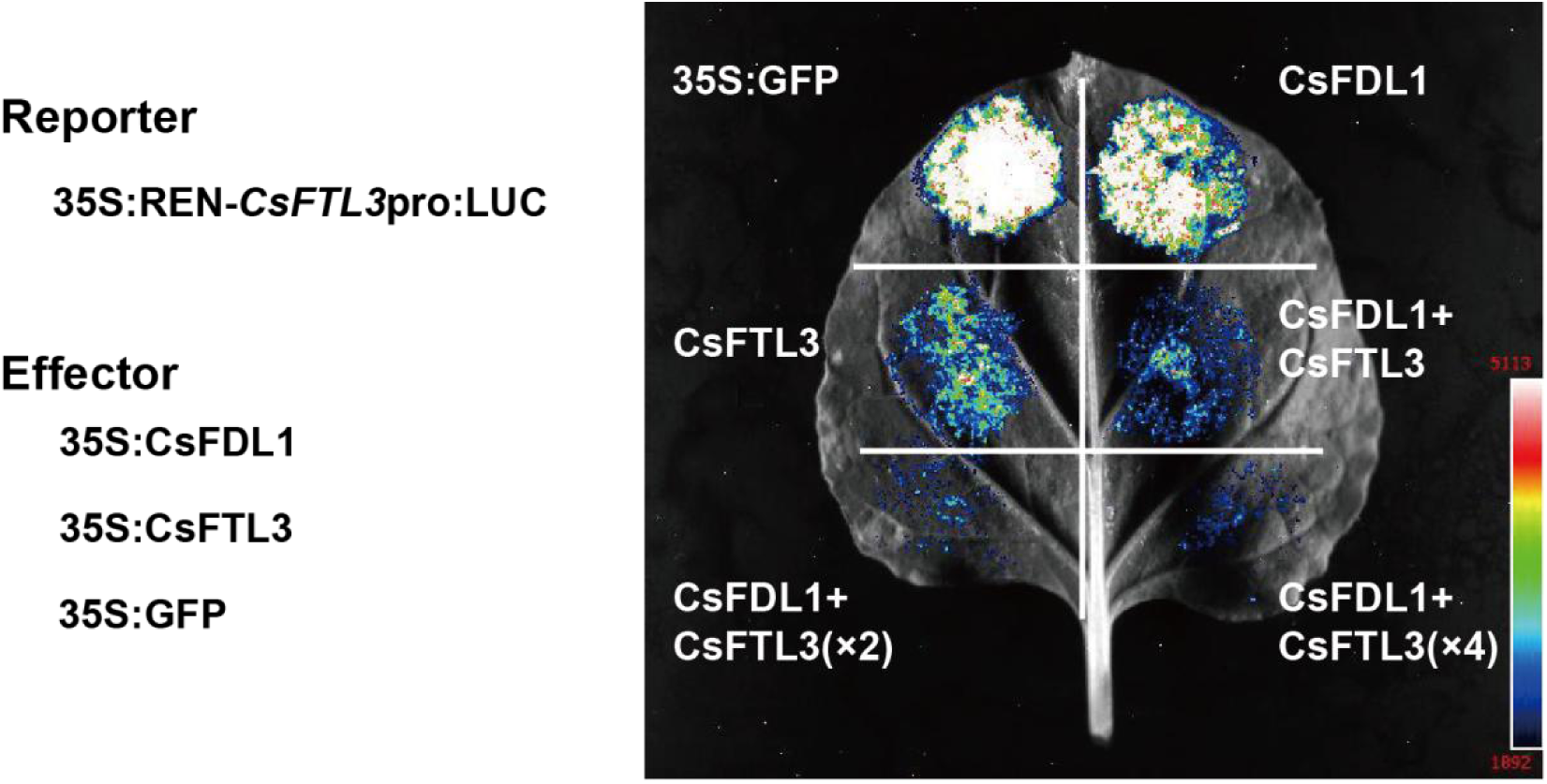
Schematic diagram of the structure of the reporter and effectors in the dual luciferase reporter system. CsFTL3-CsFDL1 complex repress the expression of *CFTL3* in tobacco cells (n >9).

**Fig.S6.**
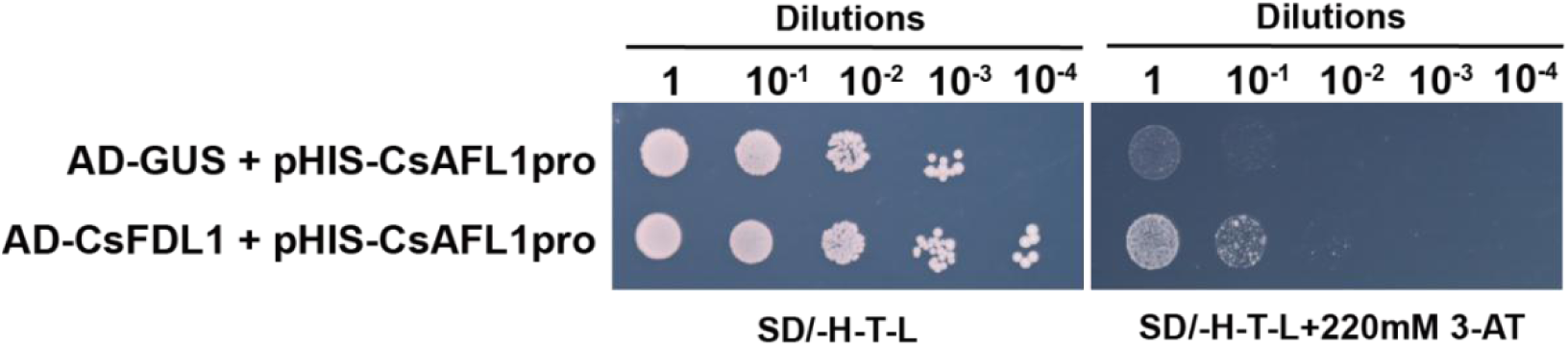
Yeast one-hybrid assay of the binding of CsFDL1 to the promoter of *CsAFL1*.

**Supplementary Table 1.**
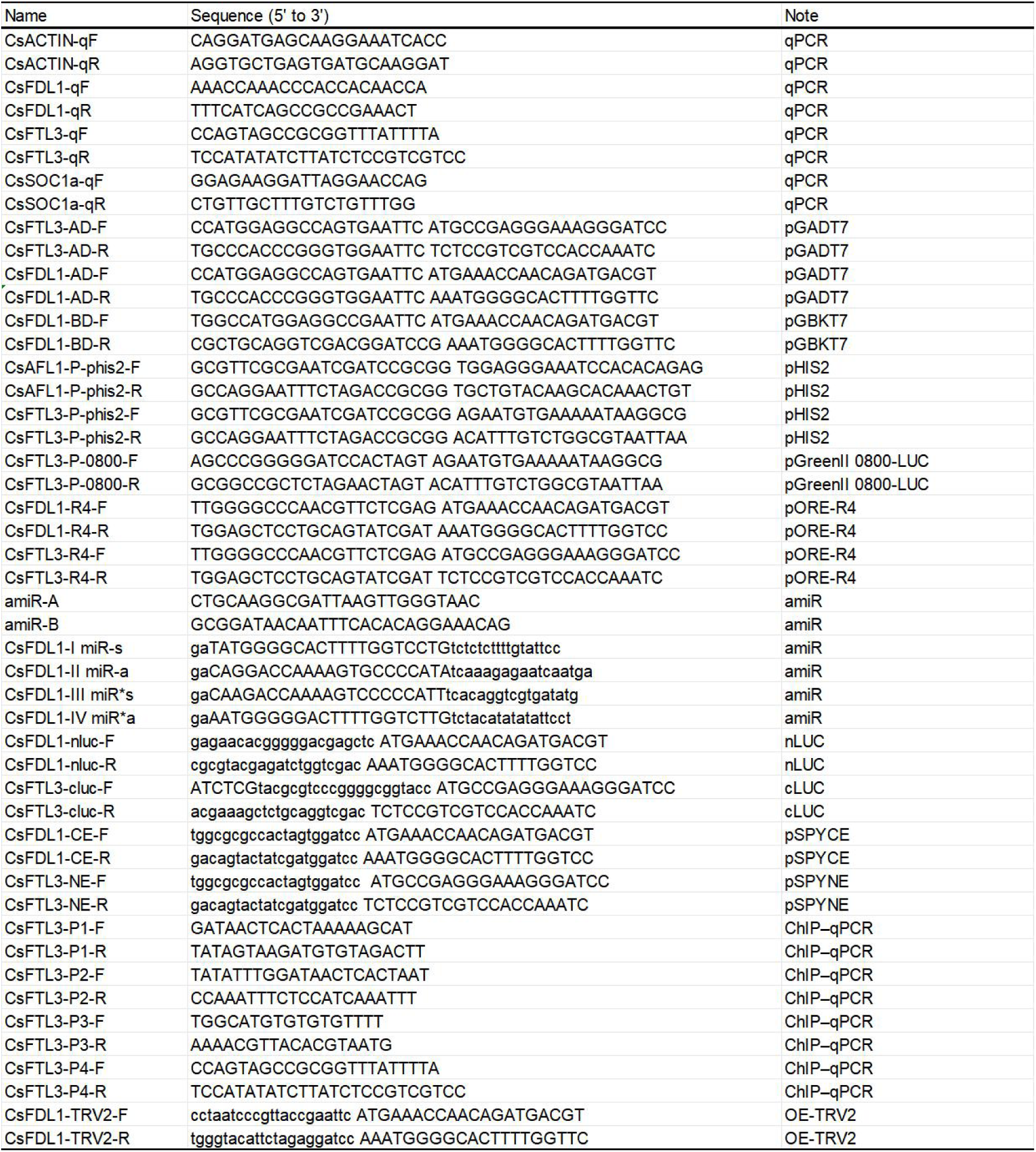
List of primers used in this study.

## References

Abe M, Kobayashi Y, Yamamoto S, et al. 2005. FD, a bZIP protein mediating signals from the floral pathway integrator FT at the shoot apex. Science 309, 1052–1056.

Andrés F, Coupland G. 2012. The genetic basis of flowering responses to seasonal cues. Nature Reviews Genetics 13, 627–639.

Balasubramanian S, Sureshkumar S, Lempe J, Weigel D. 2006. Potent induction of Arabidopsis thaliana flowering by elevated growth temperature. PLoS Genetics 2, e106.

Cheng H, Zhang J, Zhang Y, et al. 2023. The Cm14-3-3μ protein and CCT transcription factor CmNRRa delay flowering in chrysanthemum. Journal of Experimental Botany 74, 4063–4076.

Collani S, Neumann M, Yant L, Schmid M. 2019. FT modulates genome-wide DNA-binding of the bZIP transcription factor FD. Plant Physiology 180, 367–380.

Corbesier L, Vincent C, Jang S, et al. 2007. FT protein movement contributes to long-distance signaling in floral induction of Arabidopsis. Science 316, 1030–1033.

Gao H, Ding N, Wu Y, et al. 2025. Florigen activation complex forms via multifaceted assembly in Arabidopsis. Nature 648, 686–695.

Higuchi Y, Narumi T, Oda A, et al. 2013. The gated induction system of a systemic floral inhibitor, antiflorigen, determines obligate short-day flowering in chrysanthemums. Proceedings of the National Academy of Sciences of the United States of America 110, 17137–17142.

Hu Q, Yin M, Gao Z, et al. 2025. Late flowering in chrysanthemum induced by low ambient temperature is mediated by FLOWERING LOCUS C-like. Journal of Experimental Botany 76, 2192–2206.

Huang Y, Xing X, Tang Y, et al. 2022. An ethylene-responsive transcription factor and a flowering locus KH domain homologue jointly modulate photoperiodic flowering in chrysanthemum. Plant Cell and Environment 45, 1442–1456.

Izawa T, Oikawa T, Sugiyama N, et al. 2002. Phytochrome mediates the external light signal to repress FT orthologs in photoperiodic flowering of rice. Genes and Development 16, 2006–2020.

Jin S, Nasim Z, Susila H, Ahn JH. 2021. Evolution and functional diversification of FLOWERING LOCUS T/TERMINAL FLOWER 1 family genes in plants. Seminars in Cell and Developmental Biology 109, 20–30.

Kojima S, Takahashi Y, Kobayashi Y, et al. 2002. Hd3a, a rice ortholog of the Arabidopsis FT gene, promotes transition to flowering downstream of Hd1 under short-day conditions. Plant and Cell Physiology 43, 1096–1105.

Lee R, Baldwin S, Kenel F, McCallum J, Macknight R. 2013. FLOWERING LOCUS T genes control onion bulb formation and flowering. Nature Communications 4, 2884.

Li C, Lin H, Dubcovsky J. 2015. Factorial combinations of protein interactions generate a multiplicity of florigen activation complexes in wheat and barley. The Plant Journal 84, 70–82.

Li P, Song A, Gao C, et al. 2015. The over-expression of a chrysanthemum WRKY transcription factor enhances aphid resistance. Plant Physiology and Biochemistry 95, 26–34.

Lin MK, Belanger H, Lee YJ, et al. 2007. FLOWERING LOCUS T protein may act as the long-distance florigenic signal in the cucurbits. The Plant Cell 19, 1488–1506.

Liu Y, Li X, Ma D, Chen Z, Wang JW, Liu H. 2018. CIB1 and CO interact to mediate CRY2-dependent regulation of flowering. EMBO Reports 19, e45762.

Lv X, Zeng X, Hu H, et al. 2021. Structural insights into the multivalent binding of the Arabidopsis FLOWERING LOCUS T promoter by the CO-NF-Y master transcription factor complex. The Plant Cell 33, 1182–1195.

Ma L, Guan Z, Wang Q, et al. 2020. Structural insights into the photoactivation of Arabidopsis CRY2. Nature Plants 6, 1432–1438.

Nakano M, Hirakawa H, Fukai E, et al. 2021. A chromosome-level genome sequence of Chrysanthemum seticuspe, a model species for hexaploid cultivated chrysanthemum. Communications Biology 4, 1167.

Nakano Y, Takase T, Takahashi S, Sumitomo K, Higuchi Y, Hisamatsu T. 2019. Chrysanthemum requires short-day repeats for anthesis: Gradual CsFTL3 induction through a feedback loop under short-day conditions. Plant Science 283, 247–255.

Oda A, Narumi T, Li T, Kando T, Higuchi Y, Sumitomo K, Fukai S, Hisamatsu T. 2012. CsFTL3, a chrysanthemum FLOWERING LOCUS T-like gene, is a key regulator of photoperiodic flowering in chrysanthemums. Journal of Experimental Botany 63, 1461–1477.

Shu Tian, Luo X, Cui B, He Y. 2025. Auto-downregulation of the florigen FT production prevents precocious flowering in plants. bioRxiv.

Song YH, Shim JS, Kinmonth-Schultz HA, Imaizumi T. 2015. Photoperiodic flowering: time measurement mechanisms in leaves. Annual Review of Plant Biology 66, 441–464.

Sun D, Zhang J, He J, et al. 2022. Whole-transcriptome profiles of Chrysanthemum seticuspe improve genome annotation and shed new light on mRNA–miRNA–lncRNA networks in ray florets and disc florets. BMC Plant Biology 22, 515.

Taoka K, Ohki I, Tsuji H, et al. 2011. 14-3-3 proteins act as intracellular receptors for rice Hd3a florigen. Nature 476, 332–335.

Tamaki S, Matsuo S, Wong HL, Yokoi S, Shimamoto K. 2007. Hd3a protein is a mobile flowering signal in rice. Science 316, 1033–1036.

Tang QY, Zhang CX. 2012. Data Processing System (DPS) software with experimental design, statistical analysis, and data mining developed for use in entomological research. Insect Science.

Tian C, Zhai L, Wang J, Zhu W, Shi C, Jiang J, Zhao K, Li F, Zhou L, Song A, Xiong G, Li S, Chen F, Chen S. 2025. CmARF3–CmTCP7 module regulates flowering time in chrysanthemum. Horticulture Research 12, uhaf095.

Tylewicz S, Tsuji H, Miskolczi P, Petterle A, Azeez A, Jonsson K, Shimamoto K, Bhalerao RP. 2015. Dual role of tree florigen activation complex component FD in photoperiodic growth control and adaptive response pathways. Proceedings of the National Academy of Sciences of the United States of America 112, 3140–3145.

Wigge PA, Kim MC, Jaeger KE, et al. 2005. Integration of spatial and temporal information during floral induction in Arabidopsis. Science 309, 1056–1059.

Freytes SN, Canelo M, Cerdán PD. 2021. Regulation of Flowering Time: When and Where?. Current Opinion in Plant Biology 63, 102049.

